# Tegmental atrophy in isolated REM sleep behaviour disorder: *Ex vivo* MRI–informed *in vivo* imaging

**DOI:** 10.1101/2025.10.29.685226

**Authors:** Masakazu Hirose, Kenji Yoshinaga, Yoshifumi Mori, Atsushi Shima, Nobukatsu Sawamoto, Noritaka Wakasugi, Hirohiko Imai, Masahiko Watanabe, Goichi Beck, Yuta Kajiyama, Ujiakira Nishiike, Hideki Mochizuki, Kazuya Kawabata, Keita Hiraga, Tomohiko Nakamura, Masahisa Katsuno, Hirohisa Watanabe, Taku Hatano, Koji Kamagata, Nobutaka Hattori, Akira Nishida, Yohei Mukai, Eiko N. Minakawa, Yuji Takahashi, Ryosuke Takahashi, Takashi Hanakawa, Japan-Parkinson’s Disease Progressive Markers Initiative (J-PPMI) cohort

## Abstract

Isolated rapid eye movement (REM) sleep behaviour disorder (iRBD) is an early-stage synucleinopathy characterized by brainstem pathology. In rodents, the pontine tegmentum contains an REM sleep centre, the sublaterodorsal nucleus (SLD), which expresses corticotropin-releasing hormone binding protein (CRHBP). While the involvement of brainstem pathophysiology is thus implicated in iRBD, its solid evidence remains scarce in humans due to the difficulty in identifying small brainstem nuclei with conventional MRI technology alone. Here, we aimed to detect tegmental atrophy in iRBD with voxel-based morphometry (VBM) analysis combined with a novel human brainstem atlas. Structural MRIs from 98 patients with iRBD and 114 controls were analysed to investigate grey matter volume (GMV) using VBM. Our unique approach involved detailed assessments of the VBM results, guided by a high-resolution MRI-based atlas of the human brainstem. This brainstem atlas was founded on *ex vivo* MRI of 10 *postmortem* human specimens. We validated it with CRHBP immunostaining, which aided in identifying putative REM sleep-regulating nuclei in humans. We applied this brainstem atlas to identify atrophy in specific brainstem regions in iRBD and correlate their volumes with clinical measures, including autonomic functions. VBM revealed a focal cluster of grey matter atrophy in the dorsal pontine tegmentum of iRBD patients, including the laterodorsal tegmental nucleus, ventral part (LDTgV) and the pedunculopontine tegmental nucleus (PTg). Our atlas-based analysis confirmed the LDTgV as the site of most conspicuous atrophy, revealing a significant volume reduction in iRBD patients compared to controls with a moderate effect size (Cohen’s d = 0.46, Bonferroni-corrected *p* = 0.019). Furthermore, greater atrophy in the LDTgV and the PTg was associated with more severe autonomic dysfunction as measured by Scales for Outcomes in Parkinson’s Disease-Autonomic dysfunction (SCOPA-AUT) scores (partial *r* = -0.237, *p* = 0.019 and partial *r* = -0.236, *p* = 0.019, respectively). Histological analysis confirmed that the LDTgV is selectively enriched with CRHBP-positive neurons, a putative marker for REM sleep-on neurons. We provided novel evidence for the involvement of LDTgV, the putative human homolog of the murine SLD, in iRBD. The present findings advance our understanding of the neuroanatomical basis of iRBD and will contribute to the development of early biomarkers for _α_-synucleinopathies.

## Introduction

Isolated rapid eye movement (REM) sleep behaviour disorder (iRBD) is characterized by dream-enacting behaviours and REM sleep without atonia. Cohort studies of iRBD have shown that 60–76 % of patients develop _α_-synucleinopathy—Parkinson’s disease (PD), dementia with Lewy bodies (DLB) or multiple system atrophy (MSA)—within 10 years after diagnosis ^1–3^. This high conversion rate has led iRBD to be widely recognized as one of the strongest prodromal markers of _α_-synucleinopathy ^4,5^. A common and salient feature of this prodromal phase is early autonomic dysfunction, often preceding motor and cognitive decline and predicting conversion to _α_-synucleinopathy ^1,6–8^. Nevertheless, the specific neurodegenerative processes underlying iRBD and its associated features remain unclear.

The brainstem, particularly the ponto-mesencephalic tegmentum, plays a pivotal role in regulating REM sleep, as inferred from animal models ^9–11^. This region controls sleep states through a “flip-flop” mechanism involving neurons that are specifically active during REM sleep (REM-on) and those that suppress REM sleep (REM-off) ^12,13^. In rodents, these REM-on neurons are identified in the sublaterodorsal tegmental nucleus (SLD), the pedunculopontine tegmental nucleus (PTg) and the laterodorsal tegmental nucleus (LDT). These neurons interact with medulla oblongata neurons, regulating REM sleep and atonia. Among these, the SLD, also known as the subLDT ^14^, appears to be the critical region in regulating REM sleep. A recent study has identified glutamatergic neurons expressing corticotropin-releasing hormone binding protein (CRHBP) in the murine SLD, and these neurons are crucial in generating muscle atonia during REM sleep^15^. Notably, in *postmortem* investigations of human patients with PD and RBD, the CRHBP-immunoreactive neurons in the pontine tegmentum contained pathological _α_-synuclein^15^. This seminal study for the first time connected the pontine tegmentum and the pathophysiology of RBD in both rodents and humans.

However, translating these fundamental neuroanatomical insights into neuroimaging markers from living human patients remains a substantial challenge. Although previous *in vivo* structural MRI studies have reported alterations in grey matter volume (GMV) in iRBD using voxel-based morphometry (VBM), findings have been inconsistent concerning the brainstem ^16–18^. Indeed, although a recent VBM meta-analysis consistently found GMV changes in some cortical and subcortical regions, it failed to identify a uniform pattern of GMV changes within the brainstem ^19^. Beyond these inter-study discrepancies, a hurdle is the difficulty in precisely identifying specific brainstem nuclei *in vivo*. For instance, while neuromelanin MRI enables the visualization of a small nucleus, such as the locus coeruleus (LC), ^20–24^, the precise localisation of other nuclei remains challenging.

Most importantly, the human homolog of the SLD remains under debate, with candidates including the subcoeruleus nucleus (SubC) and the ventral subdivisions of the LDT (LDTgV) ^14,25,26^. Resolving this debate requires the precise differentiation of these candidate nuclei from adjacent structures, including the separation between the dorsal (i.e., LDTg) and ventral (i.e., LDTgV) parts of the LDT complex, a task not feasible with conventional approaches. A comprehensive, high-resolution, histologically validated brainstem atlas is essential for enhancing our understanding of the neuroanatomical basis of iRBD and its relationship to _α_-synucleinopathy progression.

Here, to address brainstem abnormalities in iRBD, we combined an in-house, ultra-high-resolution atlas of the human brainstem with structural MRI analyses of 98 iRBD patients and 114 matched controls from a multi-centre cohort. We examined 10 *postmortem* human brainstems using *ex vivo* 7-T MRI, visualizing the overlap of small brainstem nuclei across the ten specimens in the standard MRI space. We validated this probabilistic brainstem atlas with histology, including CRHBP immunohistochemistry (IHC). Our primary hypothesis was that this atlas-assisted approach would reveal atrophy of specific brainstem nuclei, corresponding to the putative human homologue of the SLD, which would be overlooked with conventional MRI techniques alone. Considering the prominence of autonomic dysfunction as an early feature of iRBD, we further hypothesized that structural alterations within key brainstem regions implicated in iRBD pathophysiology would correlate with the severity of autonomic symptoms. Integrating high-resolution *ex vivo* mapping and *in vivo* structural MRI analysis allowed us to identify brainstem markers of prodromal _α_-synucleinopathy in humans and their clinical relevance.

## Materials and methods

### *In vivo* MRI analysis

#### Cohort participants and data acquisition

We analysed data from 98 patients with polysomnography (PSG)-confirmed iRBD and 114 healthy aged people. The patients with iRBD participated in the Japanese Parkinson’s Progression Markers Initiative (J-PPMI) at five hospitals in Japan: The National Center of Neurology and Psychiatry (NCNP) hospital (n = 54), Juntendo University Hospital (n = 20), Kyoto University Hospital (n = 12), The University of Osaka Hospital (n = 4), and Nagoya University Hospital (n = 8). A detailed description of the J-PPMI cohort has been previously published ^27^. In brief, the J-PPMI cohort recruited males and females aged 60 years or older diagnosed with iRBD. The diagnosis of iRBD followed the International Classification of Sleep Disorders criteria (3rd revision) ^28^, including confirmation of REM sleep without atonia using PSG. Exclusion criteria included neurological or psychiatric disorders other than iRBD, a Geriatric Depression Scale (GDS)-15 score of 10 or higher, or a State-Trait Anxiety Inventory (STAI) Form Y-1 score of 65 or higher. The healthy aged control group (HA) consisted of individuals recruited from the NCNP Hospital (n = 83) and Kyoto University Hospital (n = 31). The study protocol was approved by the local ethics committees (A2014-127 in NCNP, 16-154 in Juntendo, C1059-6 in Kyoto, 14454 in Osaka, and 2015-0169 in Nagoya).

The patients with RBD completed a battery of clinical evaluations: The Japanese version of the Montreal Cognitive Assessment (MoCA-J) for cognitive function, Scales for Outcomes in Parkinson’s Disease-Autonomic Dysfunction (SCOPA-AUT) for autonomic symptoms, Movement Disorder Society-Sponsored Revision of the Unified Parkinson’s Disease Rating Scale (MDS-UPDRS) for motor symptoms, and REM Sleep Behavior Disorder Screening Questionnaire-Japanese (RBDSQ-J) for screening RBD symptoms. For evaluating cognitive functions in the control HA group, the Mini-Mental State Examination (MMSE) score was available, which was subsequently converted to estimated MoCA scores using a proposed algorithm ^29^. All participants underwent 3-dimensional (3D) T1-weighted structural MRI scanning at a magnetic field strength of 3 T. The imaging protocol was standardized across all sites, primarily using a magnetization-prepared rapid acquisition with gradient echo (MPRAGE) sequence with an isotropic resolution of 1 mm (Supplementary Table 1). Scanning parameters were optimized for each scanner while maintaining comparable image quality.

#### Voxel-based morphometry (VBM) analysis

We performed image processing and VBM analysis with SPM12 (https://www.fil.ion.ucl.ac.uk/spm/software/spm12/) implemented on MATLAB R2022 (MathWorks, Natick, MA). The processing pipeline included bias field correction and tissue-class segmentation using the standard tissue probability maps. Spatial normalisation to the Montreal Neurological Institute (MNI152) space utilized a template for East Asian brains, with the diffeomorphic anatomical registration through the exponentiated Lie algebra algorithm. Jacobian modulation was applied to GMV images to preserve absolute tissue volumes, followed by smoothing with a 6-mm full-width at half-maximum Gaussian kernel. To mitigate the effects of different MRI scanners, ComBat harmonization was applied at each voxel of the spatially normalized GMV images, modelling age, sex, total intracranial volume (TIV) and disease label as covariates of interest, while accounting for site effects. The harmonisation procedure was validated by confirming a substantial reduction in between-site variance after the ComBat correction (see Supplementary Fig. 1, Supplementary Table 2).

The site–harmonized GMV images were fed into VBM analysis with age, gender and TIV as covariates. According to our hypothesis, we limited our search volumes to the brainstem within a mask created from the MNI152 atlas. A threshold for statistical significance was set at corrected *p* < 0.05 for multiple comparisons using threshold-free cluster enhancement (TFCE).

### *Ex vivo* MRI analysis

#### *Ex vivo* MRI Data Acquisition

To locate the brainstem nuclei of interest, we acquired *ex vivo* MRI data from 10 human brainstem specimens (one male and nine females; mean age, 91.0 ± 6.3 years; range, 77–97 years) that were free from known conditions affecting brainstem morphology (see Supplementary Table 3 for details). This study protocol was approved by the Kyoto University Ethics Committee (R3074).

All specimens underwent a standardized preparation protocol. After perfusion fixation and extraction, human brain specimens were maintained in a 10% neutral buffered formalin at 4°C for at least one month, followed by more than 4 rounds of replacement with phosphate-buffered saline (PBS). Then the brainstem was dissected at the level of the mammillary bodies and immersed in PBS containing 10 mM gadoteridol for about 10 days to enhance the signal-to-noise ratio. Before scanning, each brainstem was fixed with sponges inside a plastic container filled with Fluorinert FC-43 (3M, Maplewood, MN, USA) and underwent a degassing procedure for half an hour to eliminate air bubbles that could affect image quality.

All *ex vivo* imaging was performed on a 7 Tesla MRI system for small animals (Bruker BioSpec 70/20 USR), equipped with a circular-polarized volume coil (72 mm inner diameter) for signal transmission and reception. The acquisition protocol employed a fast low-angle shot sequence (echo time/repetition time = 6.0/15.0 ms, flip angle = 60°) to achieve an isotropic resolution of 78.1 _μ_m.

#### Construction of Probabilistic Brainstem Atlas

We applied a “prefiltered rotationally invariant non-local means filter” ^30^ to enhance the signal-to-noise ratio of *ex vivo* MRIs while preserving fine anatomical details. Guided by the human brainstem atlas ^31,32^ and the 7-T WIKIBrainStem template ^33,34^, we defined criteria for a set of brainstem nuclei. A trained neuroscientist outlined seven nuclei that are recognized components of REM-sleep or arousal circuits: The PTg, the LDTg, the LDTgV, the LC, and the SubC on each side, and the midline dorsal raphe nucleus (DR) and periaqueductal grey (PAG) ^14,25,35–37^. We manually delineated volumes of interest (VOIs) on each sample’s MRI in its original high-resolution space. Another neuroscientist reviewed every contour, and disagreements were resolved by consensus. The comprehensive delineation protocol for each of the seven nuclei is provided in Supplementary Table 4. For instance, the PTg is located in the caudolateral midbrain tectum, extending from the midbrain–pontine junction, and is bordered ventrally by the superior cerebellar peduncle decussation. LDTgV is situated dorsolateral to the rostral pontine reticular formation, showing intermediate signal intensity, and bordered ventrally by the central tegmental tract (ctg), medially by the medial longitudinal fasciculus (mlf), dorsally by the LDTg, and laterally by the LC. Most nuclei presented with relatively high signal intensities compared to the low signal intensities of adjacent white matter tracts. We were unable to delineate the SLD as a distinct VOI, primarily because its precise anatomical location in humans remains a subject of ongoing debate and is not consistently defined in existing human brain atlases ^31–34^. Instead, we defined the SubC and the LDTgV separately based on established criteria as candidate regions of the SLD homologue in humans.

The *ex vivo* MRI data exhibited inter-sample variability in size, shape, and deformation, which resulted from the tissue fixation and imaging preparation processes. We developed and applied a dedicated image-processing pipeline to accommodate these variances and accurately transform each *ex vivo* MRI into the MNI152 space while minimising variability. The spatial normalisation of the *ex vivo* brainstem MRI data began with affine and non-linear diffeomorphic warping, referencing a brainstem tissue probability map adapted from the standard template. Optimal transformation parameters, determined by comparing results from standard normalisation packages with Advanced Normalization Tools (ANTs) and SPM12, were applied to the VOIs derived from each sample’s original high-resolution space, yielding a 0.5-mm resolution probabilistic atlas in the MNI152 space Finally, these spatially normalized VOIs were averaged across all samples, and voxels with a probabilistic distribution of 30% or greater for each nucleus were included in the present probabilistic brainstem atlas. The resulting three-dimensional spatial relationship of these nuclei within the probabilistic atlas is illustrated in Fig. 1B.

**Figure 1.**
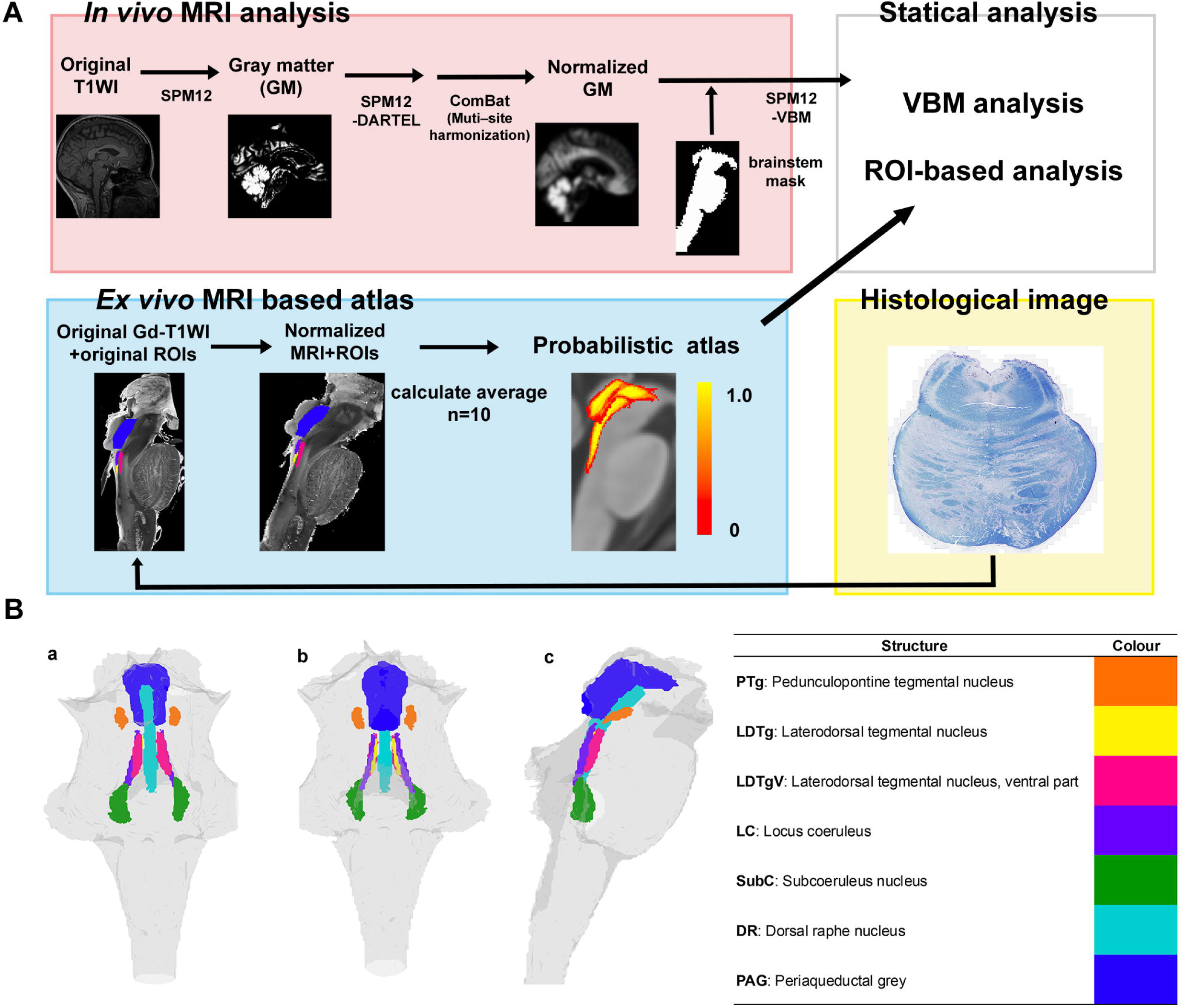
Schematic of the study workflow and the resulting three-dimensional probabilistic brainstem atlas. (A) A schematic diagram of the study: (1) in vivo MRI analysis, including grey matter extraction, DARTEL normalization, and ComBat harmonization; (2) ex vivo MRI-based probabilistic atlas construction from 10 postmortem brainstems; (3) statistical analyses, including voxel-based morphometry (VBM) and volume-of-interest (VOI) analysis; and (4) histological validation. (B) Three-dimensional renderings of the probabilistic brainstem atlas. The spatial arrangement of the seven delineated brainstem nuclei is presented from (a) anterior, (b) posterior and (c) lateral perspectives. The color key for each nucleus is provided on the right.

#### Atlas-Based Analyses

We performed two complementary analyses using the probabilistic brainstem atlas to identify which nuclei might be affected in iRBD. First, we examined the spatial overlap between the GMV reduction identified in the VBM analysis and each nucleus defined in the atlas. Second, we extracted GMV from each VOI. For the bilaterally delineated VOIs— namely the PTg, LDTg, LDTgV, LC and SubC —the VOIs were averaged across the left and right sides to yield a single value for the subsequent analysis. Group comparisons were performed for the seven nuclei, using a general linear model with age, sex, and TIV as covariates. The family-wise error rate was controlled at an _α_ of 0.05 using Bonferroni correction for multiple comparisons across the seven nuclei. We investigated the correlation between the GMV of the selected nuclei and clinical measures in the iRBD group. The primary clinical score of interest was the SCOPA-AUT total score, which assessed autonomic dysfunction in iRBD. As exploratory analyses, we also examined correlations with cognitive performance (MoCA-J) or motor impairment (MDS-UPDRS part III). Pearson’s partial correlation coefficients were calculated for these analyses, controlling for age, sex, and TIV. Both GMV and clinical scores were residualized by regressing out the effects of age, sex, and TIV before plotting. All analyses were two-tailed. We performed a primary, hypothesis-driven correlation analysis between GMV and the SCOPA-AUT total score, as well as additional exploratory analyses with the MoCA-J and MDS-UPDRS part III scores. For all correlation analyses, the threshold was set at *p* < 0.05.

#### Histological validation of the brainstem atlas

We performed histological validation of the brainstem nuclei that exhibited GMV reduction in iRBD. After the *ex vivo* MRI acquisition, the brainstem specimens were sectioned at a thickness of 15 _μ_m. We performed Klüver-Barrera staining to visualize the general cyto/myeloarchitectural organization of the brain. For IHC, deparaffinized sections were subjected to antigen retrieval by autoclaving for 15 minutes, followed by membrane permeabilization with 0.2% Triton-X for 15 minutes. Endogenous peroxidase activity was quenched with 0.3% hydrogen peroxide for 30 minutes. After washing with PBS, sections were incubated with primary antibodies. We used a goat polyclonal anti-human choline acetyltransferase (ChAT) antibody (1:50; E. coli–derived recombinant human ChAT isoform R [Ala2–Pro630], AF3447, R&D Systems, Minneapolis, MN, USA) as a marker for cholinergic neurons, and a rabbit polyclonal anti-human CRHBP antibody (1:100; human CRHBP amino acids 98-136, consistent with previous research^15^) as a marker for REM sleep-regulating neurons. The sections were treated with a secondary antibody (Histofine Simple Stain MAX-PO(G) for goat and MAX-PO(R) for rabbit; Nichirei Biosciences, Tokyo, Japan). The immunoreaction was visualized using 3,3’-diaminobenzidine with the VECTASTAIN ABC Kit (Vector Laboratories, Burlingame, CA, USA), followed by counterstaining with hematoxylin. Stained sections were imaged using an upright Axio Observer microscope (Carl Zeiss Microscopy GmbH, Germany) equipped with ZEN 3.8 software. The distribution patterns of CRHBP-and ChAT-immunoreactive neurons were compared with the VOIs on the *ex vivo* MRI.

To quantify the distribution of immunoreactive neurons, three representative *post mortem* specimens of the ten were analysed. The VOIs from our probabilistic brainstem atlas, corresponding to the anatomical levels of the midbrain, rostral pons, and caudal pons, were overlaid onto the digitized histological images of the three representative specimens. Specifically, we used the PTg VOI at the midbrain level, the LDTgV at the rostral pontine level, and the SubC at the mid-to-caudal pontine level. Within these atlas-defined regions, the number of ChAT- and CRHBP-positive cells was manually counted, and the area of each region was measured. The density of immunoreactive cells was then calculated and expressed as cells per square millimetre (cells/mm²). Within these atlas-defined regions, the number of ChAT- and CRHBP-positive cells was manually counted, and the area of each region was measured using QuPath v0.6.0 ^38^. Cell density was then calculated and expressed as cells per square millimetre (cells/mm²).

## Results

### Demographic and clinical characteristics

The iRBD group (n = 98; mean age 70.8±5.6 years; 78 males [79.6%]) and control HA group (n=114; mean age 68.9±5.7 years; 57 males [50.0%]) showed differences in age (*t*_(207)_ = -2.36, *p* = 0.019, two-tailed) and sex distribution (chi-square test, *p* < 0.001) (Table 1). Hence, age and sex were treated as confounding variables. The estimated MoCA-J scores ^29^ of the control HA group (24.8±2.7) showed no group-wise differences with the MoCA-J scores of the iRBD group (24.7±3.1) (*t*_(193)_ = 0.41, *p* = 0.67, two-tailed).

### VBM Analysis

The VBM analysis identified a single cluster of GMV reduction in patients with iRBD compared to HAs. This cluster was located in the midbrain and upper pontine tegmentum (peak MNI coordinate: *x* = -4, *y* = -34, *z* = -21; peak *t* = 4.01; 469 voxels; TFCE-corrected *p* < 0.05; Fig. 2). Our probabilistic brainstem atlas indicated that this cluster included the bilateral LDTgV, PTg, LDTg and SubC. A semi-quantitative assessment of spatial overlap revealed that 70.6% of the LDTgV volume was contained within the VBM cluster, representing the highest degree of overlap among all examined nuclei, followed by the PTg (45.9%) and LDTg (25.0%) (see Supplementary Fig. 2 for details).

**Figure 2.**
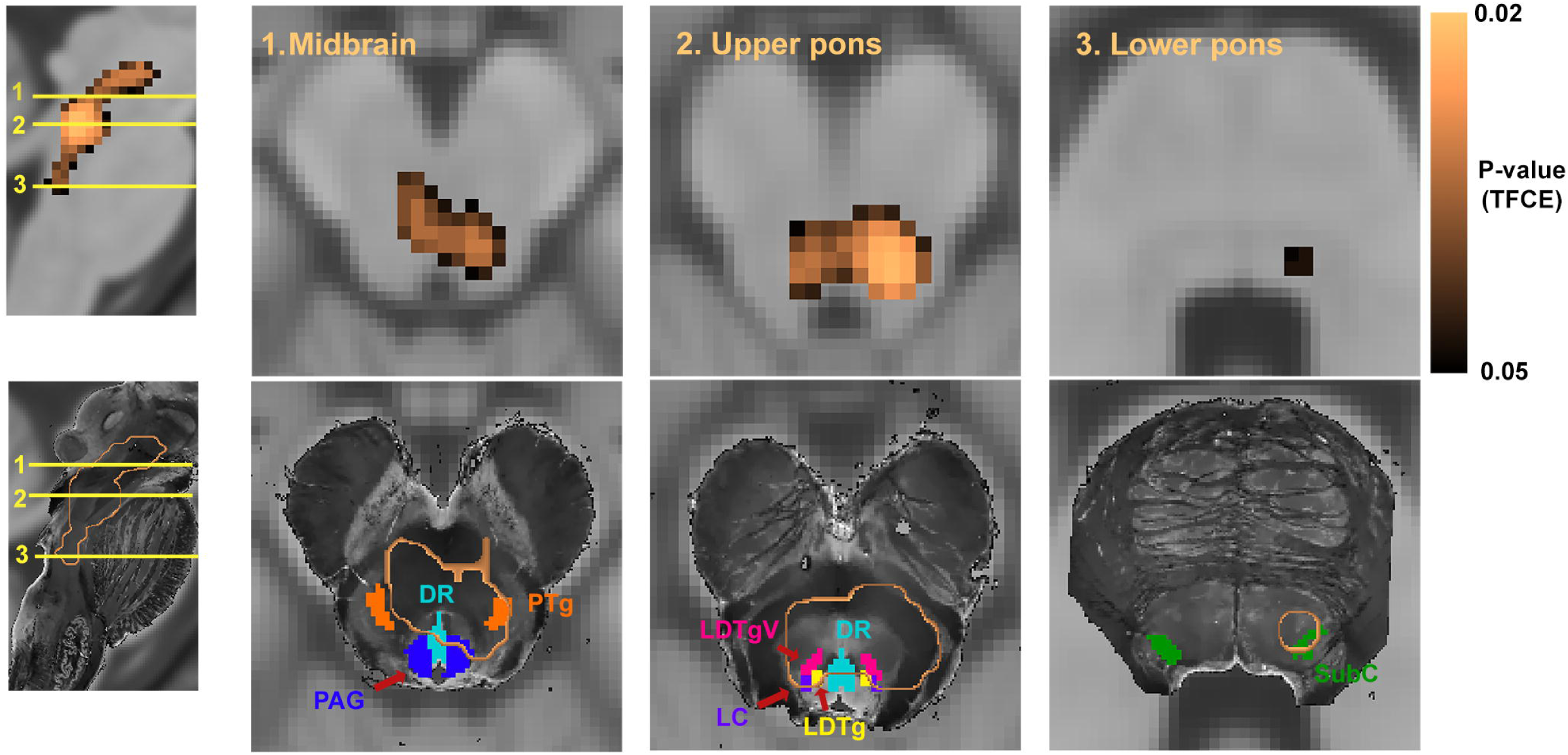
Grey matter volume (GMV) reduction in iRBD. Results of the voxel-based morphometry analysis. The top row displays a cluster of significant GMV reduction (copper) in the midbrain and upper pontine tegmentum in iRBD compared to controls (*p* < 0.05 threshold-free cluster enhancement [TFCE] corrected). The bottom row shows the VBM results (copper outlined) overlaid on the *ex vivo* brainstem atlas, demonstrating spatial overlap of the atrophy with the LDTgV, PTg, LDTg and SubC. The color bar indicates the *p*-value. Abbreviations: iRBD, isolated REM sleep behaviour disorder; LDTgV, Laterodorsal tegmental nucleus, ventral part; PTg, pedunculopontine tegmental nucleus; SubC, Subcoeruleus nucleus

### Atlas-based analysis

In the atlas-based analysis of GMV in the seven brainstem nuclei, we found significant GMV reductions in the LDTgV of patients with iRBD compared with HAs (mean difference = 0.61, 95% confidence interval [CI] [0.26, 0.97], Bonferroni-corrected *p* = 0.019, Cohen’s d = 0.46; Fig. 3), after controlling for confounding variables. Similarly, the PTg showed reduced GMV in iRBD patients compared to HAs (mean difference = 0.36, 95% CI [0.14, 0.58], Bonferroni-corrected *p* = 0.049, Cohen’s d = 0.43; Fig. 3). By contrast, GMV in the other five nuclei—the LDTg, LC, SubC, DR and PAG—did not differ significantly between the iRBD and HA groups (Bonferroni corrected *p* > 0.05).

**Figure 3.**
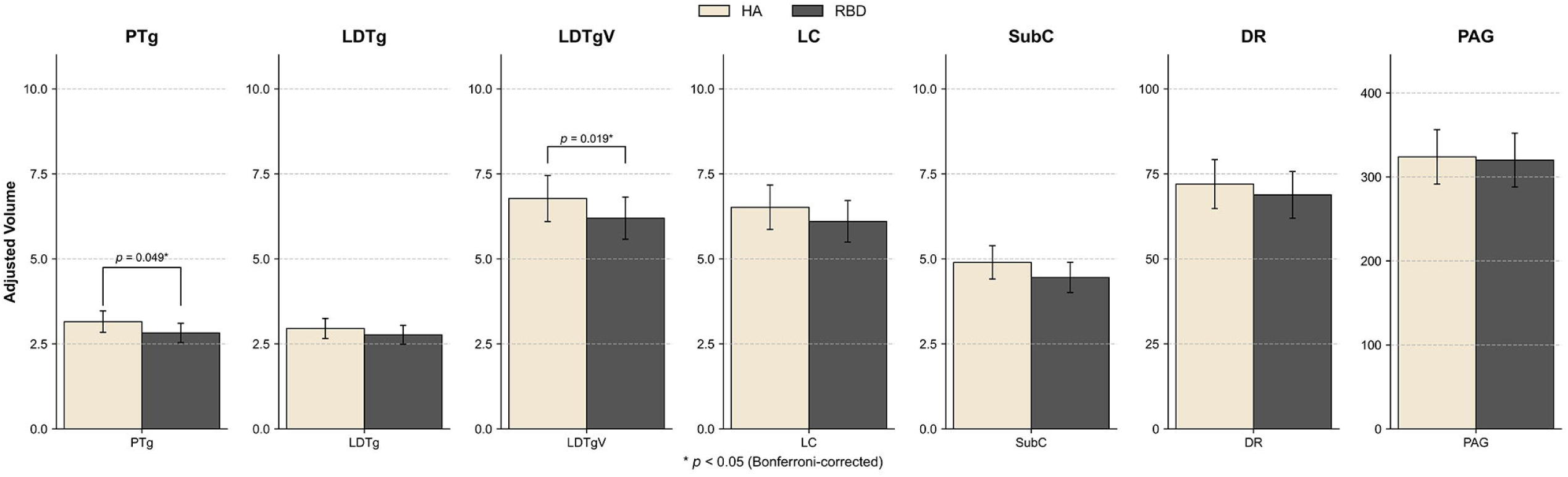
VOI-based grey matter volume analysis of brainstem nuclei. The comparison of adjusted grey matter volume in the seven brainstem nuclei between iRBD (dark grey) and controls (light beige). After Bonferroni correction for multiple comparisons, significant volume reduction was identified only in the LDTgV (*p* = 0.019) and the PTg (*p* = 0.049). Error bars represent 95% confidence intervals. Abbreviations: iRBD, isolated REM sleep behaviour disorder; LDTgV, Laterodorsal tegmental nucleus, ventral part; PTg, Pedunculopontine tegmental nucleus

### Correlation analysis

We investigated the correlation between clinical scores and atrophy in the LDTgV and the PTg, which showed structural alterations in iRBD. We focused on the correlates of autonomic dysfunction, as measured by SCOPA-AUT, after controlling for age, sex, and TIV. Partial correlation analysis revealed that smaller GMV in the LDTgV and the PTg was associated with higher standardized SCOPA-AUT scores (LDTgV: partial *r* = -0.237; 95% CI [-0.416, -0.040], *p* = 0.019, Fig. 4A; PTg: partial *r* = -0.236; 95% CI [-0.415, -0.040], *p* = 0.019, Fig. 4B), suggesting that greater atrophy in these pontine tegmental nuclei was associated with more severe autonomic dysfunction in patients with iRBD. We also conducted exploratory correlation analyses to investigate the relationship between the volumes of the LDTgV and PTg and other clinical domains, including cognitive function (MoCA-J) and motor impairment (MDS-UPDRS-III). None of these scores yielded statistically significant correlations with the LDTgV and PTg (all *p* > 0.05, uncorrected).

**Figure 4.**
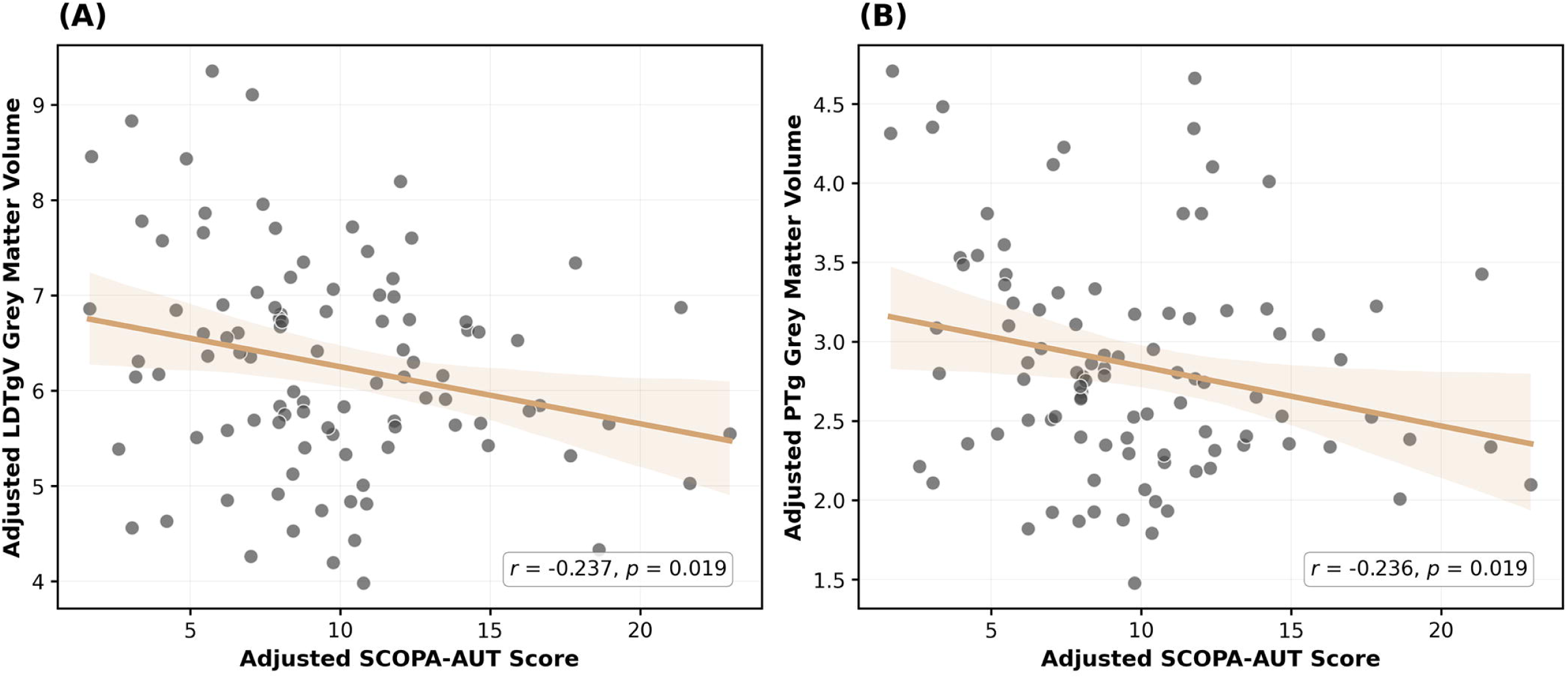
Correlation between brainstem nuclei volume and autonomic dysfunction. Scatter plots of the relationship between adjusted grey matter volume (GMV) and the Scales for Outcomes in Parkinson’s Disease-Autonomic (SCOPA-AUT) scores in patients with iRBD (n = 98). GMV and SCOPA-AUT scores were adjusted for age, sex, and total intracranial volume (TIV). Values plotted are residuals after adjustment for these covariates. (A) A significant negative correlation was observed between the LDTgV volume and SCOPA-AUT scores (*r* = -0.237, *p* = 0.019). (B) A significant negative correlation was also observed between the PTg volume and SCOPA-AUT scores (*r* = -0.236, *p* = 0.019). Solid lines represent the line of best fit; shaded areas represent the 95% confidence intervals. Abbreviations: iRBD, isolated REM sleep behaviour disorder; LDTgV, Laterodorsal tegmental nucleus, ventral part; PTg, Pedunculopontine tegmental nucleus

### Histological validation

Using *postmortem* human brainstem specimens, we conducted histological examinations of LDTgV, PTg, LDTg, LC, SubC, and DR. The Klüver-Barrera staining delineated the general cyto- and myelo-architecture. It supported that the cellular arrangements and fibre tract patterns (Fig. 5, Klüver-Barrera columns) corresponded well with the anatomical segmentations of the DR, PTg, LDTg, LDTgV, LC and SubC between the *ex vivo* MRI and histology (Fig. 5A) left columns). Consistent with established neuroanatomy, ChAT-positive cholinergic neurons were observed within the LDTg and LDTgV (Fig. 5A middle row, ChAT column), as well as in the PTg (Fig. 5A top row, ChAT column). IHC for CRHBP revealed that CRHBP-immunoreactive neurons were predominantly distributed within the LDTgV (Fig. 5A middle row, CRHBP column, arrowheads; lower row). By contrast, in the adjacent LDTg and other examined nuclei, including the DR, PTg, LC and SubC, CRHBP-immunoreactive neurons were relatively sparse or absent under the identical IHC condition. This specific pattern of CRHBP immunoreactivity within the LDTgV, when considered alongside the significant atrophy of the LDTgV observed in iRBD *in vivo,* suggests that this neuronal population is preferentially targeted by the underlying pathological process.

**Figure 5.**
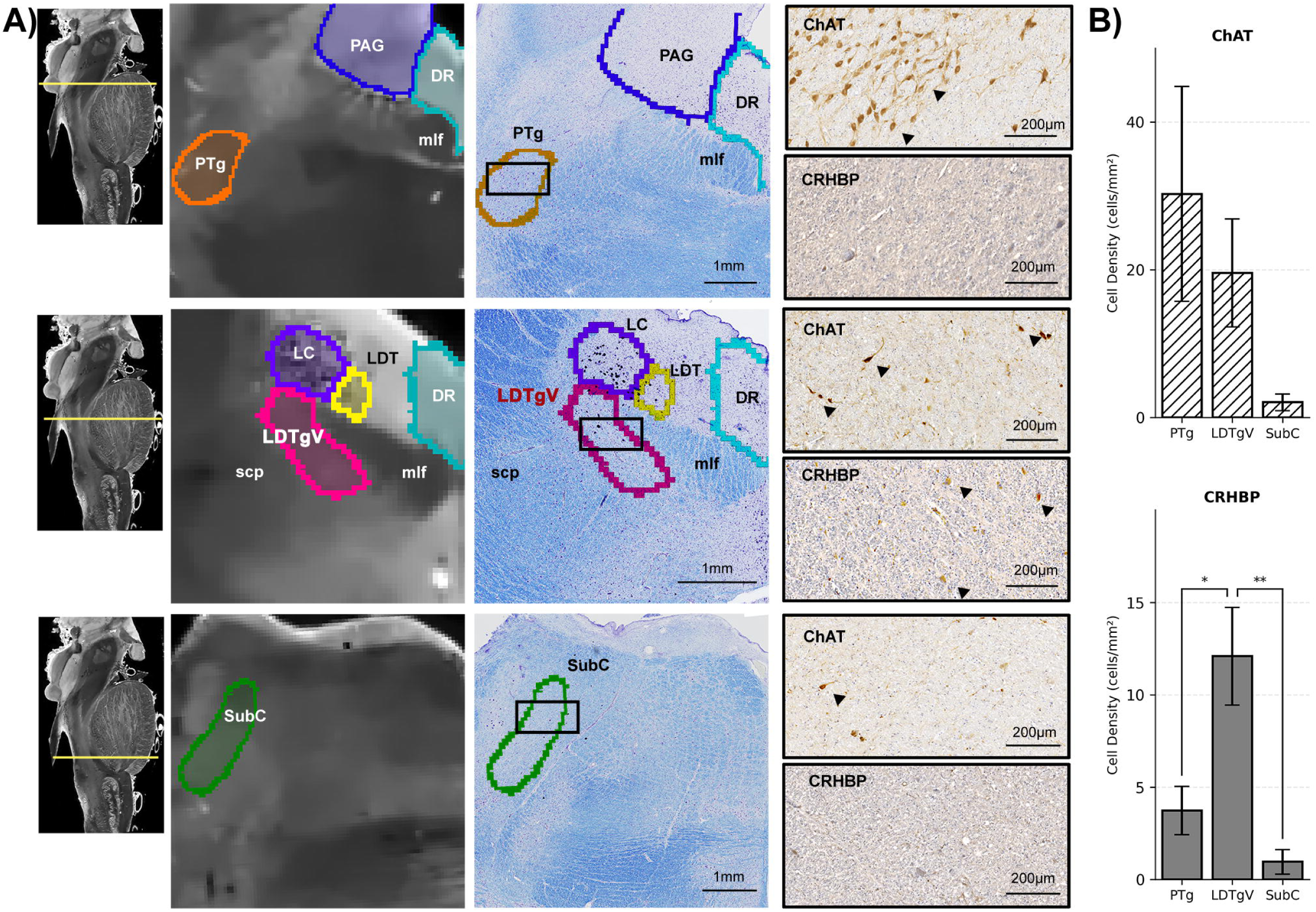
Histological validation of brainstem nuclei. **(A)** Each row corresponds to a brainstem level shown in the sagittal MRI. Columns from left to right show: (1) *ex vivo* MRI with color-coded nuclei delineations; (2) Klüver-Barrera staining for cyto- and myeloarchitecture; (3) Choline acetyltransferase (ChAT) immunohistochemistry (corresponding to the black rectangle in (2)); (4) Corticotropin-releasing hormone-binding protein (CRHBP) immunohistochemistry. The areas in (3) and (4) correspond to the black rectangle in (2). Arrows indicate immunoreactive neurons. CRHBP-positive neurons, a marker for REM sleep-on neurons, were localized to the LDTgV. **(B)** Quantitative cell density analysis of ChAT- and CRHBP-immunoreactive neurons in the PTg, LDTgV, and SubC. Cell densities were calculated from three *postmortem* human specimens (n = 3 brains). The density of CRHBP-positive neurons was markedly higher in the LDTgV compared to the other nuclei. Values are presented as mean ± standard error of the mean (SEM). Statistical comparisons were made using a one-way ANOVA followed by Tukey’s post-hoc test. For CRHBP, significant differences were found between PTg and LDTgV (* *p* < 0.05) and between LDTgV and SubC (** *p* < 0.01). ChAT-positive neurons were distributed across all three nuclei without significant differences. Abbreviations: LDTgV, Laterodorsal tegmental nucleus, ventral part; PTg, Pedunculopontine tegmental nucleus; SubC, Subcoeruleus nucleus

Quantitative analysis of the three representative specimens supported these observations, revealing distinct distribution patterns of CRHBP- and ChAT-immunoreactive neurons across the brainstem nuclei (Fig. 5B). The density of CRHBP-positive neurons was highest in the LDTgV (12.08 ± 2.64 cells/mm²), followed by the PTg (3.73 ± 1.31 cells/mm²), and was minimal in the SubC (0.96 ± 0.66 cells/mm²). ChAT-positive neurons were most densely distributed in the PTg (30.26 ± 14.53 cells/mm²) and LDTgV (19.57 ± 7.29 cells/mm²), with the lowest density in the SubC (2.06 ± 1.14 cells/mm²). This quantitative data reinforces that CRHBP-expressing neurons, implicated in regulating REM sleep, are particularly concentrated within the LDTgV, the putative human homolog of the murine SLD, located in the rostral pontine tegmentum.

## Discussion

We analysed structural MRI data from a cohort of iRBD and HAs, guided by a newly developed histology-validated human brainstem atlas based on *ex vivo* 7 T MRI. The VBM findings found GMV reductions in the midbrain and upper pontine tegmentum in iRBD. The atlas-based volumetric analysis revealed atrophy in both the LDTgV and the PTg in iRBD compared to HAs. The LDTgV exhibited the most pronounced GMV reduction among the brainstem nuclei. Our IHC analysis pinpointed the LDTgV that harbours CRHBP-expressing neurons, which have been proven to be a specific neuronal group regulating REM sleep. The human homolog of the murine SLD has been a subject of debate. Previous literature often regarded the SubC as this human equivalent ^14,25,26^; however, the supporting evidence is not solid. Another line of evidence, based on detailed studies of the rat pontomesencephalic tegmentum, suggests that the LDTgV may correspond to the murine SLD ^39,40^. Our histological examination clearly demonstrated CRHBP immunoreactivity in the LDTgV, whereas the SubC showed only sparse CRHBP immunoreactivity. Therefore, our findings are consistent with this anatomical definition, supporting the LDTgV rather than the SubC as the human homolog of SLD.

The CRHBP-expressing neurons within the human LDTgV/murine SLD are of paramount importance for REM sleep atonia^15^. Animal model studies have established that this specific neuronal population is indispensable for generating muscle atonia during REM sleep. For instance, targeted ablation of these CRHBP-positive SLD neurons drastically reduces REM sleep and impairs REM atonia in mice ^15^. The loss of the CRHBP-expressing neurons, which co-localized with _α_-synuclein pathology, was also reported in *postmortem* brainstems of PD with RBD ^15^, providing direct evidence linking the human LDTgV/SLD to _α_-synucleinopathies. The PTg, another REM-related nucleus that showed atrophy in our iRBD cohort, showed only sparse CRHBP immunoreactivity, further highlighting the unique neurochemical profile of the LDTgV. Our current findings, combined with the animal evidence for the role of CRHBP neurons in REM atonia, suggest that the human LDTgV, the putative homolog of the murine SLD, underlies the pathophysiology of iRBD.

Beyond its role in REM atonia, the LDT complex serves as an integrator of brainstem functions characterized by extensive connectivity. This aligns with evidence from human tractography studies, which show projections from the LDT complex to the thalamus, hypothalamus, basal forebrain, and frontal cortex, as well as to other brainstem nuclei, including the PTg ^41^. The LDT complex is implicated not only for REM sleep and arousal ^42^ but also for autonomic regulation via the nucleus of the solitary tract and parabrachial nuclei ^41^. Hence, the atrophy of the LDTgV likely extends its functional impact to a broader array of non-motor symptoms, including autonomic dysfunctions, which are frequently associated with iRBD. This notion is supported by a significant association between LDTgV atrophy and SCOPA-AUT scores, which aligns with a previous large-scale network study^43^. The LDTgV, with its established connections to autonomic centres and its demonstrated vulnerability in iRBD, may therefore represent a critical anatomical substrate whose degeneration is linked to both the REM sleep dysregulation and the coexisting autonomic disturbances. Identifying such specific brainstem correlates of non-motor symptoms in iRBD was critically dependent on the anatomical precision afforded by our high-resolution, histologically validated atlas, which precisely delineates the LDTgV from adjacent structures and quantifies these specific pathological changes.

Devised from the gross VBM findings, the precise localisation of brainstem atrophy to the LDTgV and PTg has helped disambiguate the existing, more varied landscape of brainstem pathology suggested for iRBD. Earlier structural MRI studies have implicated brainstem abnormalities in iRBD, though the specifically involved nuclei and the consistency of their volumetric changes remain unclear. For instance, some VBM reports indicated GMV reductions in the pontine tegmentum^44^, while others showed more diffuse GMV loss in the brainstem^43^. In contrast, one study combining DTI and VBM found microstructural alterations in the midbrain and pons using DTI, but no significant brainstem GMV reductions^45^. Indeed, a meta-analysis of VBM literature has failed to show a uniform pattern of brainstem atrophy in iRBD^19^, reflecting not only methodological differences across studies but also challenges in resolving small brainstem nuclei with standard technology. The utilisation of our newly developed brainstem atlas helped advance the understanding of the pathophysiology of RBD. This approach enabled the clear identification of atrophy in LDTgV and PTg, identifying specific neuroanatomical correlates for the VBM-detected changes. Thus, by identifying these specific nuclei, our findings provide a more anatomically precise understanding of the pathophysiology of iRBD than previous reports.

The LDTgV atrophy in our iRBD cohort may mirror the _α_-synuclein pathology previously identified in CRHBP-expressing neurons in the corresponding region of *postmortem* brains from patients with PD and RBD^15^. Together, the LDTgV may be an early target of _α_-synuclein pathology. The early and significant pathological involvement of a key REM-atonia regulating nucleus, such as the LDTgV, may therefore underpin the neuroanatomical basis for the manifestation of iRBD. Considering that not all individuals in the prodromal phase of _α_-synucleinopathies develop iRBD, the pronounced LDTgV atrophy in our iRBD cohort may indicate that this specific clinical phenotype is tightly linked to substantial pathological changes within this nucleus. It is plausible that iRBD-predominant prodromal _α_-synucleinopathy is characterized by an earlier or more severe involvement of the LDTgV and its associated circuits, compared to prodromal individuals who may present with other non-motor symptoms first. Future longitudinal studies comparing brainstem structural changes in prodromal individuals with and without iRBD are crucial to test these hypotheses regarding distinct pathophysiological trajectories.

Our study had several limitations. This was a cross-sectional study, precluding conclusions about the temporal progression of atrophy or its direct causal link to clinical conversion. Future work would benefit from longitudinal data analysis to track LDTgV and PTg atrophy and their predictive value for conversion to specific _α_-synucleinopathies. The lack of significant atrophy in REM-related nuclei other than the LDTgV and PTg does not necessarily exclude their involvement in RBD. Indeed, structural connectivity alterations involving the SubC have been reported in iRBD^46^, suggesting that different types of pathological changes may occur in different nuclei or be detectable by different modalities. Integrating multimodal neuroimaging, including DTI to assess structural connectivity and functional MRI to evaluate brainstem network activity, will be crucial. The application of our detailed brainstem atlas to investigate other neurodegenerative disorders affecting the brainstem also represents a promising path for future research. Regarding the *ex vivo* atlas, tissue fixation introduced anatomical variability that may affect boundary precision between adjacent nuclei. The probabilistic nature of our atlas, integrating 10 specimens, addresses inter-individual variations. Future development of automated segmentation methods using machine learning approaches may further improve the objectivity and reproducibility of brainstem nuclei delineation.

In conclusion, we provide robust evidence that iRBD is associated with atrophy of the LDTgV, a CRHBP-rich pontine tegmental nucleus, which we considered a human homolog of the murine SLD. This degeneration of the LDTgV, alongside the PTg, was associated with autonomic dysfunction, suggesting a contributory role of these pontine tegmental structures in the pathophysiology of iRBD. These findings enhance our understanding of the neuroanatomical basis of iRBD and highlight the LDTgV as a potential anatomical site for developing a biomarker for prodromal _α_-synucleinopathy.

## Data availability

The probabilistic brainstem atlas generated during this study will be made publicly available through a dedicated repository upon publication. The raw *ex vivo* magnetic resonance imaging data and full-resolution histological images are available from the corresponding author upon reasonable request. The de-identified individual-level *in vivo* magnetic resonance imaging and clinical data from the J-PPMI cohort are not publicly available due to privacy regulations and stipulations within the participant consent forms.

