## Supplementary Fig. 1 for "Tegmental atrophy in isolated REM sleep behaviour disorder: *Ex vivo* MRI–informed *in vivo* imaging"

**Supplementary Figure 1**: **Effect of ComBat harmonization on multi–site MRI data.**

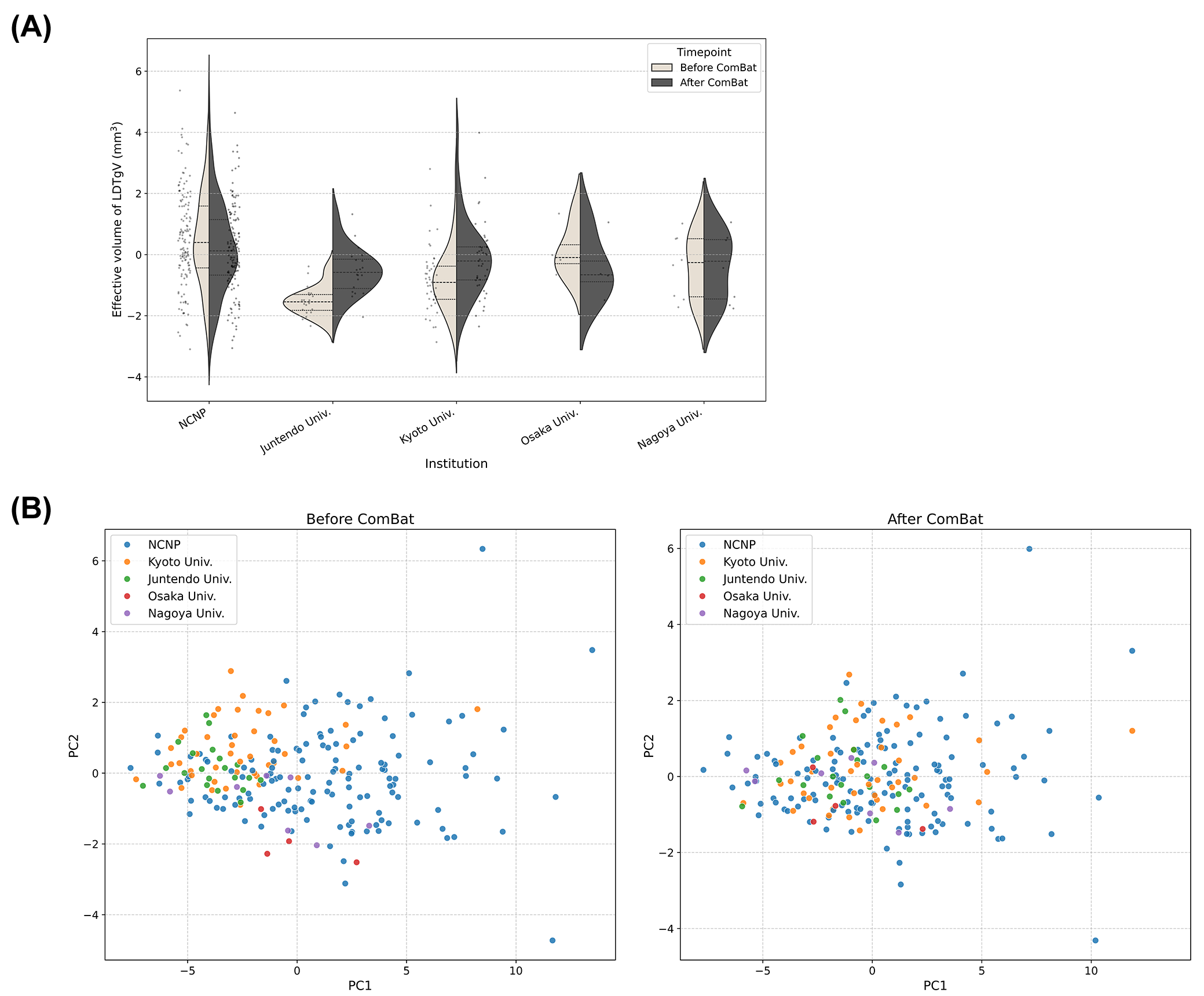

(A) A split violin plot showing the distribution of effective volume for a representative VOI (LDTgV) across the sites. The left half of each violin (beige) shows the data distribution before ComBat, and the right half (dark grey) shows the one after ComBat. Individuals’ data are shown as dots. The data distribution becomes comparable across the sites after ComBat harmonization. (B) Principal component analysis of volumes of all brainstem nuclei examined. Before ComBat (left panel), individuals’ data tended to form a cluster following the sites. The clusters were not clear after ComBat (right panel), supporting successful removal of the site effect from the data structure.

Abbreviations: NCNP, National Center of Neurology and Psychiatry; Juntendo Univ., Juntendo University; Kyoto Univ., Kyoto University; Osaka Univ., University of Osaka; Nagoya Univ., Nagoya University; PC, Principal component

**S**
