## Supplementary Fig. 2 for "Tegmental atrophy in isolated REM sleep behaviour disorder: *Ex vivo* MRI–informed *in vivo* imaging"

**Supplementary Figure 2:** Brainstem nucleus volume overlapping with the VBM atrophy cluster

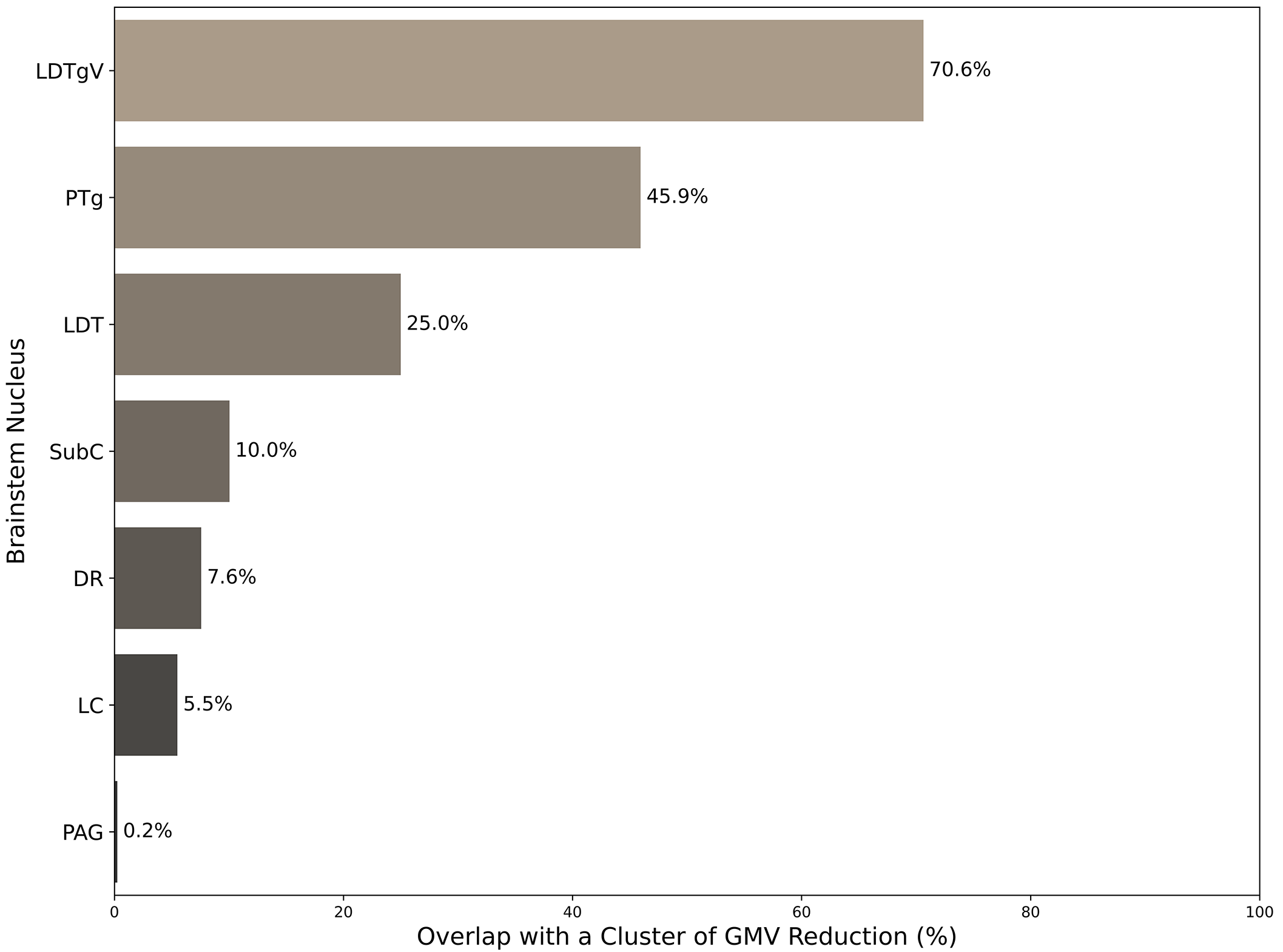

The *x*-axis represents the percentage of the atrophic volume overlapping each nucleus, divided by the total volume of each nucleus, in iRBD. Nuclei are sorted in descending order of the overlap percentage. The atrophic volume in iRBD primarily overlapped the LDTgV and PTg.

Abbreviations: LDTgV, Laterodorsal tegmental nucleus, ventral part; PTg, pedunculopontine tegmental nucleus; LDTg, Laterodorsal tegmental nucleus; SubC, Subcoeruleus nucleus; DR, Dorsal raphe nucleus; LC, Locus coeruleus; PAG, Periaqueductal grey
