## Supplementary Table 5 for "Tegmental atrophy in isolated REM sleep behaviour disorder: *Ex vivo* MRI–informed *in vivo* imaging"

**Supplementary Table 5: Demographic and clinical characteristics of participants by institution.**

|  | **HA**  **(NCNP)** | **HA**  **(Kyoto)** | **iRBD**  **(NCNP)** | **iRBD**  **(Kyoto)** | **iRBD**  **(Juntendo)** | **iRBD**  **(Osaka)** | **iRBD**  **(Nagoya)** |
| --- | --- | --- | --- | --- | --- | --- | --- |
| Age (years) | 68.9±5.8 | 68.9±5.7 | 69.6 ±5.9 | 72.2 ±4.4 | 73.3 ±5.1 | 70.0 ±4.6 | 71.9 ±6.0 |
| Sex (male: female) | 43:40 | 14:17 | 38:16 | 15:5 | 11:1 | 3:1 | 6:2 |
| MoCA-J | 24.8±2.2(estimated) | 25.0±3.8(estimated) | 24.9±3.0 | 23.3±3.5 | 25.1±1.9 | 25.5±3.3 | 25.4±4.5 |
| MMSE | 28.7±1.1 | 28.6±1.9 | NA | NA | NA | NA | NA |
| SCOPA-AUT | NA | NA | 8.1±3.7 | 12.0±6.1 | 9.8±2.9 | 13.8±4.6 | 11.9±4.4 |
| MDS-UPDRS Part III | NA | NA | 1.8±2.5 | 2.6±7.1 | 3.5±3.5 | 1.8±2.1 | 6.3±5.4 |
| RBDSQ-J | NA | NA | 7.6±3.0 | 6.7±3.1 | 6.8±1.7 | 10.8±2.5 | 7.6±3.9 |

This table provides a detailed breakdown of demographic data (age and sex) and key clinical scores (MoCA-J, MMSE, SCOPA-AUT, MDS-UPDRS Part III and RBDSQ-J) for both the healthy aged control (HA) and iRBD groups at each participating institution. All values are presented as mean ± standard deviation unless otherwise specified.

Abbreviations: NCNP, National Center of Neurology and Psychiatry; Juntendo Univ., Juntendo University; Kyoto Univ., Kyoto University; Osaka Univ., University of Osaka; Nagoya Univ., Nagoya University.
