## Supplementary Table 4 for "Tegmental atrophy in isolated REM sleep behaviour disorder: *Ex vivo* MRI–informed *in vivo* imaging"

**Supplementary Table 4:** **Delineation Criteria for Brainstem Nuclei in *Ex vivo* MRI Atlas.**

| **Delineation Criteria for Brainstem Nuclei in *Ex vivo* MRI Atlas** | |
| --- | --- |
| **Structure** | **Anatomical Location and MRI Characteristics** |
| PTg (Pedunculopontine tegmental nucleus) | Located in the reticular region of the caudolateral midbrain tectum, extending from the midbrain-pontine junction. Bordered dorsally by the isthmic reticular formation and ventrally by the superior cerebellar peduncle decussation. |
| LDTg (Laterodorsal tegmental nucleus) | Situated laterally in the pontine central grey, extending from the midbrain-pontine boundary to the mid-pons. Presents as a mildly hyperintense structure adjacent to LC. Bordered by the mlf and LDTgV ventrally, surrounded by the central grey matter. |
| LDTgV(Laterodorsal tegmental nucleus, ventral part) | Positioned dorsolateral to the rostral pontine reticular formation. Shows intermediate signal intensity (higher than the surrounding pontine reticular region, lower than the central grey). Bordered by the PnO and the ctg ventrally, the mlf medially, the LDT dorsally, and the LC laterally |
| LC (Locus coeruleus) | Located laterally in the pontine tegmentum throughout the pons. Identified by characteristic "pepper-and-salt" appearance due to neuromelanin. Appears as a round structure axially. Surrounded by PnO, LDTgV, LDT, and superior cerebellar peduncle. |
| SubC (Subcoeruleus nucleus) | Presents as a faint hyperintense area in the lateral lower pons, caudal to LC. Extends ventrally in pontine reticular formation around Mo5N medially, terminating at the rostral end of the facial nucleus (7N). |
| DR (Dorsal raphe nucleus) | Located ventral to the cerebral aqueduct, appearing as an elongated hyperintense structure extending axially. Bounded ventrolaterally by the mlf. The lateral boundary with PAG is defined by three lines extending to the cerebral aqueduct: caudally, a line from the dorsolateral angle of the mlf to the midline of the cerebral aqueduct; at mid-level, a line from the lateral margin of the trochlear nucleus (4N); and rostrally, a line from the lateral margin of the oculomotor nucleus (3N) |
| PAG (Periaqueductal grey) | High-signal grey matter structure surrounding the cerebral aqueduct in the midbrain. Extends from the caudal end of the posterior commissure rostrally to the superior medullary velum caudally. Bounded dorsally by superior and inferior colliculi, ventrally by DR, with boundaries defined by lines from mlf and cranial nerve nuclei to the cerebral aqueduct. |
| Abbreviations: ctg (Central Tegmental Tract), mlf (Medial Longitudinal Fasciculus), Mo5N (Motor Nucleus of the Trigeminal Nerve), PnO (Pontine Nucleus Oralis). | |

This table summarizes the specific anatomical landmarks and magnetic resonance imaging (MRI) signal characteristics used to manually delineate the seven brainstem nuclei of interest (PTg, LDTg, LDTgV, LC, SubC, DR, and PAG) on high-resolution 7-Tesla *ex vivo* T1-weighted images. Abbreviations used within the table are defined in the footnote.
