## Supplementary Table 3 for "Tegmental atrophy in isolated REM sleep behaviour disorder: *Ex vivo* MRI–informed *in vivo* imaging"

**Supplementary Table 3: Details of human brainstem specimens used for *ex vivo* MRI.**

| **Case** | **Age of death** | **Sex** | **Cause of death** |
| --- | --- | --- | --- |
| 1 | 94 | Female | Chronic kidney disease |
| 2 | 97 | Female | Aspiration pneumonia |
| 3 | 87 | Male | Aspiration pneumonia |
| 4 | 77 | Female | Ischemic heart disease |
| 5 | 89 | Female | Sepsis |
| 6 | 97 | Female | Senility |
| 7 | 97 | Female | Senility |
| 8 | 87 | Female | Strangulated ileus |
| 9 | 82 | Female | Acute myocardial infarction |
| 10 | 94 | Female | Cardioembolic stroke |

This table provides the demographic and specimen characteristics for the 10 *postmortem* human brainstem samples used for the high-resolution *ex vivo* MRI analysis.
