## Supplementary Table 2 for "Tegmental atrophy in isolated REM sleep behaviour disorder: *Ex vivo* MRI–informed *in vivo* imaging"

**Supplementary Table 2: Assessment of site effect reduction by ComBat harmonization**

| **VOI** | **F (Before ComBat)** | **F (After ComBat)** | **η² (Before ComBat)** | **η² (After ComBat)** |
| --- | --- | --- | --- | --- |
| **LDTgV** | 14.86 | 1.81 | 0.22 | 0.03 |
| **LDT** | 11.43 | 1.34 | 0.18 | 0.03 |
| **LC** | 10.96 | 1.55 | 0.17 | 0.03 |
| **PTg** | 24.06 | 3.06 | 0.32 | 0.06 |
| **SubC** | 12.15 | 1.97 | 0.19 | 0.04 |
| **PAG** | 24.55 | 2.18 | 0.32 | 0.04 |
| **DR** | 21.40 | 2.08 | 0.29 | 0.04 |

This table provides quantitative metrics assessing the magnitude of the site effect on grey matter volume for each brainstem nucleus before and after ComBat harmonization. The magnitude of the site effect is reported using the F-statistic (F) and partial eta-squared (η²) from a one-way ANOVA. ComBat reduced the site effects in all nuclei. Data from the left and right sides were averaged, except for the midline structures (PAG and DR). Abbreviations: Pedunculopontine tegmental nucleus (PTg), Laterodorsal tegmental nucleus (LDTg), Laterodorsal tegmental nucleus, ventral part (LDTgV), Locus coeruleus (LC), Subcoeruleus nucleus (SubC), Dorsal raphe nucleus (DR), Periaqueductal grey (PAG)
