## Supplementary Table 1 for "Tegmental atrophy in isolated REM sleep behaviour disorder: *Ex vivo* MRI–informed *in vivo* imaging"

**Supplementary Table 1: MRI acquisition parameters at each institution.**

| **Sites** | **Scanner** | **Sequence** | **TR (ms)** | **TE (ms)** | **TI (ms)** | **FA (degree)** | **Resolution (mm)** |
| --- | --- | --- | --- | --- | --- | --- | --- |
| NCNP | Siemens Verio Dot | MPRAGE | 1900 | 2.5 | 900 | 9 | 1.0×1.0×1.0 |
| Juntendo | Philips Achieva | MPRAGE | 15 | 3.4 | 1022 | 10 | 1.0×1.0×1.0 |
| Kyoto | Siemens Skyra | MPRAGE | 1900 | 2.6 | 900 | 9 | 0.9×0.9×0.9 |
| Osaka | GE Discovery MR 750 | IR-SPGR | 8.2 | 3.2 | 400 | 11 | 1.2×1.2×1.2 |
| Nagoya | Siemens Verio | MPRAGE | 2500 | 2.5 | 900 | 8 | 1.0×1.0×1.0 |

Detailed imaging parameters for the 3-dimensional T1-weighted imaging (3DT1WI) sequences used at each site. Information listed includes scanner manufacturer, sequence name, repetition time (TR), echo time (TE), inversion time (TI), flip angle (FA), and resolution.

Abbreviations: NCNP, National Center of Neurology and Psychiatry; Juntendo Univ., Juntendo University; Kyoto Univ., Kyoto University; Osaka Univ., University of Osaka; Nagoya Univ., Nagoya University.
